# A gnotobiotic mouse model with divergent equol-producing phenotype: potential for determining microbial-driven health impacts of soy isoflavone daidzein

**DOI:** 10.1101/2024.03.01.582818

**Authors:** Lindsay M. Leonard, Abigayle M. R. Simpson, Shiyu Li, Lavanya Reddivari, Tzu-Wen L. Cross

## Abstract

The implications of soy consumption on human health have been a subject of debate, largely due to the mixed evidence regarding its benefits and potential risks. The variability in responses to soy has been partly attributed to differences in the metabolism of soy isoflavones, compounds with structural similarities to estrogen. Approximately one-third of humans possess gut bacteria capable of converting soy isoflavone daidzein into equol, a metabolite produced exclusively by gut microbiota with significant estrogenic potency. In contrast, lab-raised rodents are efficient equol producers, except for those raised germ-free. This discrepancy raises concerns about the applicability of traditional rodent models to humans. Herein, we designed a gnotobiotic mouse model to differentiate between equol producers and non-producers by introducing synthetic bacterial communities with and without the capacity into female and male germ-free mice. These gnotobiotic mice display equol-producing phenotypes consistent with the capacity of the gut microbiota received. Our findings confirm the model’s efficacy in mimicking human equol production capacity, offering a promising tool for future studies to explore the relationship between endogenous equol production and health outcomes like cardiometabolic and fertility. This approach aims to refine dietary guidelines by considering individual microbiome differences.

## Introduction

The benefits of consuming soy for human health remain largely inconclusive and controversial. For example, soy consumption has been related to poor sperm quality, likely due to the estrogenic activity of soy isoflavones [1,2]. On the contrary, for health challenges related to estrogen deficiency, such as those related to menopause, soy supplementation has been shown to sometimes be beneficial [3,4]. Soy isoflavones are plant-derived phytoestrogens that have structural similarities to mammalian-synthesized estrogen and have binding capacity to estrogen receptors (ER)-alpha and -beta to act as estrogen agonists or antagonists [5]. However, inconsistent outcomes in clinical human feeding trials lead to the hypothesis that soy-derived benefit in estrogen deficiency is dependent on a person’s ability to produce (*S*)-equol (hereafter denoted as equol) [6,7]. Equol is an exclusively microbial-produced metabolite of one of the soy isoflavones, daidzein, with the highest estrogenic potency among all soy isoflavones and their metabolites [8]. An interesting phenomenon that likely contributes to the interpersonal variations in clinical outcomes of soy consumption is that only ∼30-50% of humans possess equol-producing bacteria within their gut microbiome to convert daidzein to equol [9]. In contrast, lab-raised rodents are efficient equol-producers except those that are raised germ-free [10]. Therefore, soy-feeding studies using conventionally raised rodent models may only be relevant to half of the human population at best. Developing a proper negative control (i.e., a model with a gut microbiome that cannot produce equol) is critical to discern the impact of soy consumption based on equol-producing status.

Exogenous supplementation of equol that was synthesized through *in vitro* bacterial fermentation has been tested to determine its potential benefits in humans, particularly those that do not produce equol themselves. A high level of oral equol supplementation appears to have some success in reducing cardiovascular risks and bone resorption [11–13]. However, the pharmacokinetics of oral equol supplementation differs significantly from endogenous production within the gastrointestinal tract and has been reported to provide little to no cardiometabolic benefits in non-equol-producing humans [14–16]. In fact, a recent study showed an opposing effect that dietary supplementation of equol exacerbated metabolic dysfunction in high-fat diet-induced obese mice by suppressing physical activity and energy expenditure and causing hyperglycemia, hyperinsulinemia, and hypoleptinemia [17]. Therefore, intestinal-produced equol likely exerts a distinct physiological role than exogenous supplementation of equol.

Germ-free rodents do not produce equol due to the absence of intestinal bacteria required to metabolize daidzein into equol [10]. However, living in a sterile environment is associated with various developmental and physiological abnormalities, making germ-free rodents a poor control when examining physiological and metabolic outcomes of bacterial-produced equol. Gnotobiotic mice (i.e., germ-free mice colonized with a known microbial community), on the other hand, provide the potential to serve as an appropriate negative control as non-equol producers. Bowey et al. have demonstrated this concept by colonizing gnotobiotic rats with a human microbiome of a poor equol producer to create a non-equol-producing rodent [18]. While this approach has translational relevance to human health, several downsides exist, including 1) reproducibility is low when colonization of a community relies on a human fecal sample and 2) the ecological dynamics of a complex human microbial community when encountering a novel equol-producing species is likely highly individualized due to the large interpersonal variations of the gut microbiome [19]. An alternative approach to creating a rodent model to test the endogenously synthesized equol status is to create a gnotobiotic rodent model using synthetic bacterial communities that are designed and assembled *in vitro* with precision before inoculating germ-free mice [20–22]. The benefits of using synthetic bacterial communities are greater community stability and reproducibility of colonization. A couple of attempts have been made involving the synthetic community to create equol-producing and non-equol-producing rodent models. One study compares a synthetic community that consists of eight mouse-derived bacterial strains as the non-equol-producing “control” to a conventional microbiome that produces equol to study the benefits of equol production in ApoE-null mice on reproductive health [23]. While this method utilizes a synthetic microbiome, the comparison to a complex microbiome introduces confounding variables that are not accounted for. Further, mouse-derived bacterial strains may have less relevance to human health. Another study bred gnotobiotic rats colonized with a synthetic community built with human-derived bacterial strains called the simplified human microbiota (SIHUMI), which cannot produce equol [24]. Once rats reached adulthood at 12 weeks of age, an equol-producing strain *Slackia isoflavoniconvertens* DSM 22006 was introduced to create equol-producing rats. Although much better controlled than comparing a synthetic microbiota to a complex microbiota where multiple confounding factors exist, this approach cannot be used to study the developmental or transgenerational impact of equol production as the breeders remain to be non-equol producers.

Herein, we aimed to develop a gnotobiotic mouse model colonized with synthetic microbiota from human-derived bacterial strains with distinct equol-producing capacity at the time of inoculation. We hypothesize that when two communities differ by only the presence of one bacterial strain that is capable of producing equol, the corresponding equol phenotype will be present when inoculated into germ-free mice. The development of a model system is essential to understand the impact of microbial metabolites of nutrients on human health and contribute to the development of novel strategies to maximize the nutritional efficacy of soy foods.

## Materials and Methods

### Design of Synthetic Bacterial Communities

To design these synthetic bacterial communities with disparate equol-producing capacity, we first selected a total of 10 strains that (1) are commonly found in the human gut, (2) have not been reported with equol-producing capability, and (3) were used previously in synthetic bacterial communities to colonize rodent gut [25–27]. These 10 strains were used as the “core” microbiota without equol-producing capacity (Equol(-)): *Bacteroides caccae*, *Bacteroides thetaiotaomicron*, *Bacteroides uniformis*, *Roseburia intestinalis*, *Faecalibacterium duncaniae*, *Agathobacter rectalis*, *Coprococcus comes*, *Akkermansia muciniphila*, *Providencia stuartii*, and *Collinsella aerofaciens*. To create the equol-producing (Equol(+)) community, an equol-producing strain of *Adlercreutzia equolifaciens* was added to the Equol(-) community (**Table 1**). Each strain was cultured individually and screened for equol-producing capacity in two ways: (1) conducting National Center for Biotechnology Information (NCBI) nucleotide Basic Local Alignment Search Tool (BLAST) searches against the three genes known to be involved in equol production: *dzr*, *ddr*, and *tdr* (sequences from the whole genome of the equol-producing strain *A. equolifaciens* DSMZ 19450, GenBank Accession no.: GCA_000478885.1) [28], and (2) *in vitro* culture to confirm presence and absence of equol-producing capacity before pooling to form communities.

**Table 1.**
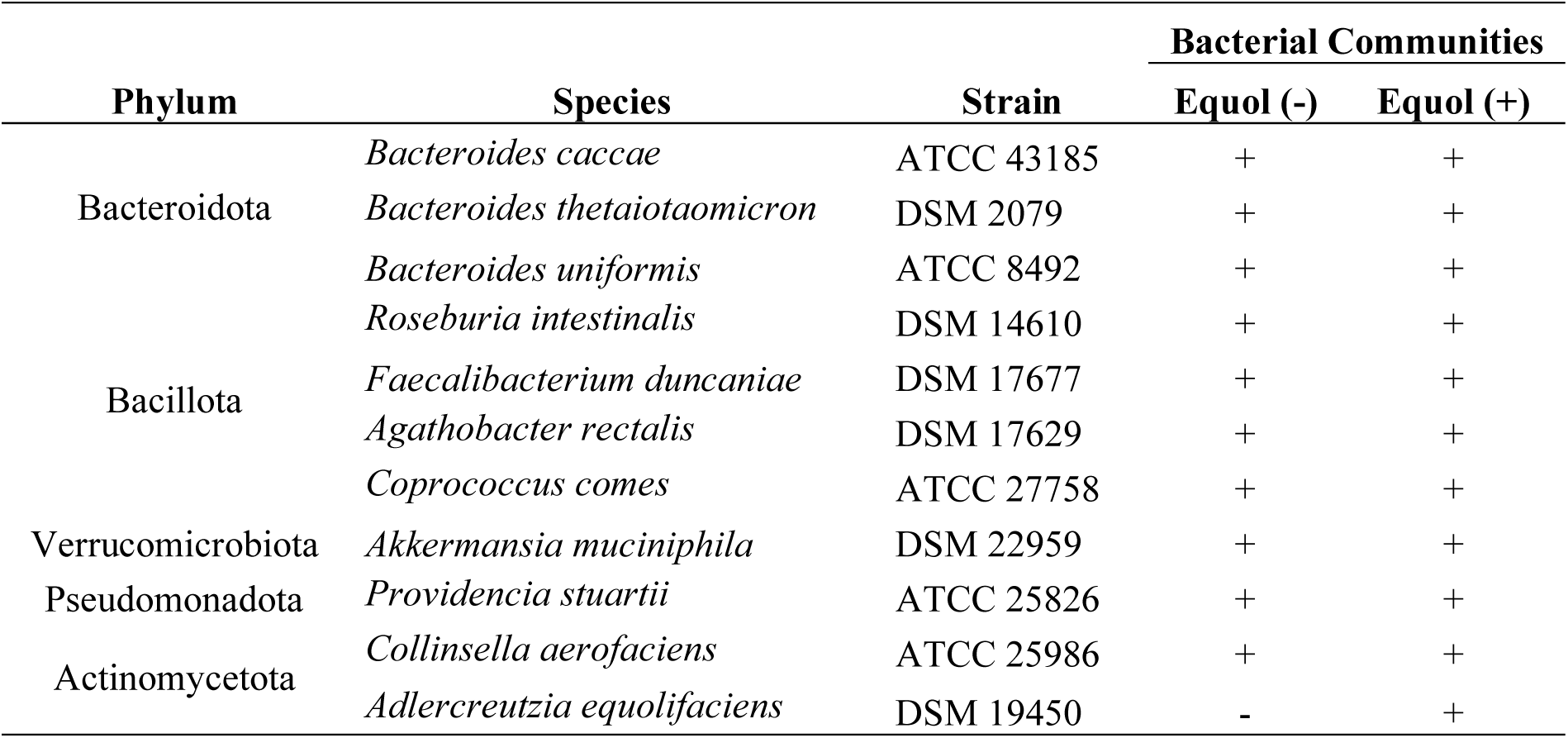
Bacterial strains included in the equol-producing (Equol(+)) and non-equol producing (Equol(-)) communities.

All strains were acquired from either the German Collection of Microorganisms and Cell Cultures (DSMZ, Braunschweig, Germany) or the American Type Culture Collection (ATCC, Manassas, VA) and revived as recommended by the ATCC or DSMZ. The culture media recipes and incubation times used for this study are designated in **Supplemental Tables 1**. All strains were grown anaerobically in an anaerobic chamber (Vinyl Anaerobic Chamber Type B, Coy Laboratory Products Inc., Grass Lake, MI) at 37°C. The identity of each strain was confirmed via 16S rRNA gene sequencing using the primer pairs 27F (5’-AGAGTTTGATCCTGGCTCAG-3’) and 1492R (5’-TACGGYTACCTTGTTACGACTT-3’) [29–31] with the Invitrogen Platinum II Taq Hot-Start DNA Polymerase kit (Cat. No. 14966001, Invitrogen, Waltham, MA). The 16S rRNA gene amplicons were sequenced by GENEWIZ (Azenta Life Sciences, South Plainfield, NJ). The taxonomy of the cleaned nucleotide sequences was then determined using the NCBI nucleotide BLAST.

Following the confirmation of the purity, identity, and equol-producing capabilities, each bacterial strain was grown separately and then pooled into the inoculants that can be used to colonize mice. Strains were first grown on agar, picked into their respective broth, and incubated. The inoculants were made by pooling each strain in equal optical density (OD_600_) at an approximate of 1.0. Two bacterial strains were not able to reach the OD_600_ of 1.0 (*A. equolifaciens* [OD_600_: 0.16], *Providencia stuartii* [OD_600_: 0.15]), so they were added into the inoculants without dilution. Using an equal volume of each strain, inoculants of the Equol(-) and Equol(+) communities were prepared and supplemented with 20% glycerol in sterile glass Balch-type tubes, and crimp-sealed, and stored at -80°C. All culture media used in this process was prepared with the oxygen indicator, resazurin (Cat. No. AC189900010, ACROS Organics, Waltham, MA) to ensure anaerobic conditions.

### Gnotobiotic mouse study

All animal protocols were approved by the Purdue Animal Care and Use Committee. Female and male germ-free C57BL/6 mice were bred and maintained at the Purdue Gnotobiotic Facility and fed Teklad Global 19% protein extruded rodent diet (sterilizable 2019S, Envigo, Indianapolis, IN) before starting this study. Mice of 4-6 weeks of age were removed from the breeding isolators and housed in ventilated cages (Sentry SPP™ Mouse, Allentown, LLC, Allentown, NJ) for the duration of the experimentation. Mice were placed on a semi-purified diet formulated based on AIN-93G formulation but adjusted to contain fermentable fibers and supplemented with 1.5 g/kg daidzein (Cat. No. D-2946, LC Laboratories, Woburn, MA). The composition of the diet is shown in **Table 2**. This diet was sterilized through double gamma irradiation at 10-20 kGy (Research Diets, New Brunswick, NJ). Food and water were provided *ad libitum* throughout the study. Mice were allowed to adjust to this diet for two weeks before bacterial inoculation. At 6-8 weeks of age, mice were colonized with one of the synthetic bacterial communities (Equol(-) or Equol(+)) through oral gavage. Inoculants were thawed on ice and remained anaerobic during this process. Immediately before colonization, fresh fecal pellets were collected to confirm the sterility of the mice before bacterial colonization. Four weeks after colonization, mice were fasted for 4 hours and euthanized using carbon dioxide asphyxiation. Cecal contents and serum were collected and flash-frozen in liquid nitrogen. Serum samples were collected from blood acquired through cardiac puncture after euthanasia and allowed to clot at room temperature for approximately 45 minutes, then centrifuged at 2,000 x g for 10 minutes at 4°C. All samples were stored at -80°C until analysis.

**Table 2.** Composition of study diet. The diet for this study was formulated based on the AIN-93G diet and was modified to be supplemented with 1.5 g/kg daidzein (LC Laboratories, D-2946) and made to contain fermentable fibers. The cellulose in the AIN-93 diet was replaced with an equal mixture by weight of the following fermentable fibers: inulin, short-chain fructo-oligosaccharides (scFOS), beta-glucan, pectin, and glucomannan. Additional alterations include using mineral acid casein instead of lactic casein, increasing to a 1.5X vitamin mix, and using corn oil in place of soybean oil. This diet was prepared by Research Diets (Research Diets, New Brunswick, NJ).

| Macronutrients | g % | kcal % |
| --- | --- | --- |
| Protein | 19% | 20% |
| Carbohydrate | 67% | 70% |
| Fat | 4% | 10% |

| Ingredient | g/kg | kcal/kg |
| --- | --- | --- |
| Casein- Mineral Acid | 190.18 | 761 |
| L-Cystine | 2.85 | 11 |
| Corn Starch | 476.59 | 1907 |
| Maltodextrin 10 | 118.86 | 475 |
| Sucrose | 60.67 | 242 |
| Inulin | 9.51 | 14.3 |
| short-chain fructo-oligosaccharides (scFOS) (93%) | 9.51 | 0.0 |
| Beta-Glucan (75.9%) | 9.51 | 5 |
| Pectin (80%) | 9.51 | 0 |
| Glucomannan NOW (98%) | 9.51 | 0 |
| Corn Oil | 23.77 | 214 |
| Lard | 19.02 | 171 |
| Mineral Mix S10026 | 9.51 | 0 |
| DiCalcium Phosphate | 12.36 | 0 |
| Calcium Carbonate | 5.23 | 0 |
| Potassium Citrate 1 H2O | 15.69 | 0 |
| Vitamin Mix V10001 | 14.26 | 57 |
| Choline Bitartrate | 1.90 | 0 |
| FD&C Red Dye #40 | 0.05 | 0 |
| Daidzein | 1.50 | 0 |

### Quantification of daidzein and equol

The equol-producing capability of each bacterial strain included in the synthetic bacterial communities was confirmed before colonizing the mice. At the conclusion of the mouse study, the concentrations of the daidzein and equol were also quantified in the serum collected from the mice to determine the equol-producing status of the mice. Each bacterial strain was grown in culture media supplemented with 100 μM daidzein (Cat. No. D-2946, LC Laboratories, Woburn, MA). The concentrations of equol and daidzein in the culture media were quantified using an Ultivo Triple Quadrupole LC-MS/MS (Model G6465A, Agilent Technologies, Santa Clara, CA) after a double ethyl acetate extraction [32–34]. An internal standard, 4-hydroxybenzophenone (4-HBP) (Cat. No. H20202, Sigma-Aldrich, St. Louis, MO), was added to each sample before extractions. For each sample, 1 ml of culture media was mixed with 6 ml of HPLC grade ethyl acetate, vortexed for 60 seconds, and centrifuged at 3,220 x g for 5 minutes at room temperature. This extraction was repeated, and the solvent phase was removed to another tube. Samples were dried using nitrogen and re-suspended in HPLC grade methanol and filtered through a 4mm PVFD membrane 0.45 μM filter unit (Cat. No. SLHV004SL, Millipore Sigma, St. Louis, MO) before analysis. Serum collected from gnotobiotic mice colonized with the Equol(-) and Equol(+) communities were analyzed for levels of daidzein and equol [35]. From each mouse, 100 μl of serum was mixed with 100 μl of acetate buffer (0.2 mol/L, pH 5.0) containing 100 units of ß-glucuronidase/aryl sulfatase (Cat. No. 1041140002, Sigma-Aldrich, St. Louis, MO) and 50 μM of the internal standard 4-HBP (Cat. No. H20202, Sigma-Aldrich, St. Louis, MO). Samples were incubated for 15 hours at 37°C, and then vortexed with 400 μl of HPLC-grade methanol and sonicated for 5 minutes. The samples were then centrifuged at 5,000 x g for 5 minutes at 4°C. The supernatant was filtered through a 4mm PVFD membrane 0.45 μM filter (Cat. No. SLHV004SL, Millipore Sigma, St. Louis, MO) to eliminate proteins. Samples were subjected to the same LC-MS/MS procedure as for bacterial culture extractions. Serum from germ-free mice was used as a negative control for equol. Standard curves were established by spiking germ-free mouse serum with a gradient of known concentrations of daidzein and equol ranging from 0.05 μM to 50 μM.

### Microbiota Analysis

The gut microbiota composition of the gnotobiotic mice was determined through 16S rRNA gene sequencing of the cecal content. The inoculants used to colonize the mice were also sequenced. DNA was first isolated from the cecal contents and inoculants using a phenol-chloroform method[36]. Briefly, homogenization of the samples was achieved by adding extraction buffer (200 mM Tris (pH 8.0), 200 mM NaCl, 20 mM EDTA), 20% SDS, and phenol:chloroform:isoamyl alcohol (25:24:1, pH 7.9) (Cat. No. 15593-049, Invitrogen, Waltham, MA) and bead beating with 0.1-mm diameter zirconia-silicate beads (Cat. No. 11079101z, BioSpec Products, Bartlesville, OK) and a single 3.2mm stainless steel bead (Cat. No. 11079132ss, BioSpec Products, Bartlesville, OK) using a Mini-Beadbeater-96 (Cat. No. 1001, BioSpec Products, Bartlesville, OK). The homogenates were centrifuged at 7,200 x g for 3 minutes at 4°C, and the aqueous layer was removed. DNA was precipitated using isopropanol and sodium acetate (3M, pH 5.2). The DNA pellets were rinsed with 100% ethanol, speed vacuumed using a Vacufuge plus (Cat. No. 022820001, Eppendorf, Hamburg, Germany), and resuspended in T_10_E_1_ buffer (10mM Tris, 1mM EDTA, pH8). The QIAquick 96 PCR purification kit (Cat. No. 28183, Qiagen, Hilden, Germany) was used to further purify the DNA. Samples were quantified using a Qubit Flex Fluorometer (Cat. No. Q3326, Invitrogen, Waltham, MA) using the Qubit broad range dsDNA kit (Cat. No. Q32853, Invitrogen, Waltham, MA). The hypervariable 4 (V4) region of the 16S rRNA gene was amplified [37]. Briefly, PCR was carried out using the KAPA HiFi HotStart Ready Mix (Cat. No. 7958935001, Roche, Basel, Switzerland). The amplified products were purified using the QIAquick PCR purification kit and underwent size selection using 1.5% low-melt agarose gels and Gel DNA Recovery Kit (Cat. No. D4002, Zymo Research, Irvine, CA). A final pool was made after quantification of each sample using the Qubit high-sensitivity dsDNA kit (Cat. No. Q32851, Cat. No. Invitrogen, Waltham, MA). Sequencing of the pool was performed using the Illumina MiSeq platform to generate 2×250bp reads by the Purdue Genomics Core. Reads were analyzed using the Quantitative Insights into Microbial Ecology version 2 (QIIME2) pipeline. Sequences were trimmed and filtered using the DADA2 software package [38]. Taxonomy was assigned to reads using a custom reference database that contained only the 11 bacterial species included in the communities.

### Strain-Specific qPCRs

To confirm the presence and quantity of the two strains that were challenging to detect through 16S rRNA gene sequencing, strain-specific quantitative PCRs (qPCRs) were performed using the DNA isolated from the cecal contents of each mouse and the bacterial inoculants. Primer sequences used to target *A. equolifaciens* (*ddr* gene, Forward: 5’-CTCGAYCTSGTSTACAACGT-3’, Reverse: 5’-GARTTGCAGCGRATKCCGAA-3’) and *F. duncaniae* (Forward: 5’-TGCCCCCGGGTGGTTCT-3’, Reverse: 5’-CGTTATTCAAAGCCCCGTTATCAA-3’) have been previously described [26,39]. Primer specificity was confirmed by testing for PCR amplification of primers against all strains used in the synthetic community. qPCR was performed using Powerup™ SYBR™ Green Master Mix (Cat. No. A25743, Applied Biosystems, Waltham, MA) following the manufacturer’s directions on a QuantStudio 7 Flex Real-Time PCR System (Cat. No. 4485701, Applied Biosystems, Waltham, MA). Amplification data and melt curves were analyzed using the QuantStudio Real-Time PCR software v1.7.1. Standard curves were established to quantify the bacterial load of each strain in the samples. Briefly, each bacterial culture was quantified through serial dilution and agar spread plating after entry into the stationary phase to determine the colony-forming units per ml of culture media (CFU/ml). DNA was isolated from the primary dilution tube and then serially diluted ten-fold to create ten standards.

### Cecal Short Chain Fatty Acids

Short-chain fatty acids (SCFAs) were quantified from the cecal contents of mice colonized with Equol(+) and Equol(-) communities using a published method [40]. Briefly, cecal contents were weighed and transferred to tubes containing 1.2 g of zirconia-silicate beads (Cat. No. 11079101z, BioSpec Products, Bartlesville, OK). Cecal contents were homogenized using a bullet blender and a vortex in 1 mL of 0.5% phosphoric acid per 100 mg of sample. The supernatant was mixed with an equal volume of ethyl acetate containing 0.14 mL heptanoic acid/L as an internal standard. This mixture was vortexed for 5 minutes before centrifugation at 17000 x g at 4°C for 10 minutes. The ethyl acetate phase was recovered and stored at -80°C. An Agilent 7890A gas chromatograph (GC-FID 7890A, Santa Clara, CA, USA) with a fused silica capillary column (Nukon SUPELCO No: 40369-03A, Bellefonte, PA, USA) was used to quantify the SCFA concentration. Peak areas for all SCFAs were recorded, corrected for extraction efficiency and sample volume variability using the internal standard heptanoic acid, and quantified using a standard curve.

### Statistical Analysis

Data was analyzed and graphed in GraphPad Prism version 9.4.1 (GraphPad Software, San Diego, CA). Outliers for daidzein and equol concentration in the serum were identified and removed using the robust regression and outlier removal (ROUT) method. For daidzein and equol serum quantification and qPCR data, normality was assessed using a Shapiro-Wilk test (alpha=0.05). Variables that passed the Shapiro-Wilk test were analyzed using an unpaired t-test or a one-way analysis of variance (ANOVA) followed by Tukey’s multiple comparisons test. For variables that did not pass the Shapiro-Wilk test, a Mann-Whitney or a Kruskal-Wallis test followed by Dunn’s multiple comparison test was used. For the microbiota data obtained through 16S rRNA gene Illumina sequencing, statistical differences in beta diversity were tested using permutational multivariate analysis of variance (PERMANOVA) in QIIME2. Permutational multivariate analysis of dispersions (PERMDISP) was also performed in QIIME2 to determine significant differences in group variances. Distance matrices were graphed as principal coordinates analysis (PCoA) plots using the package qiime2R via RStudio version 4.1.2 [41]. Ellipses presented on these graphs were calculated based on a multi-variable t distribution with a radius of 0.95 around the center of the data for each group using the stat_ellipses function in ggplot through R. Data are presented as the mean ± standard error of the mean (SEM). P values below 0.05 were considered statistically significant. Statistical significance is indicated as follows: *=p < 0.05; **=p < 0.01; ***=p < 0.001, ****=p < 0.0001.

## Results

### Body weight, mesenteric fat, and gonadal fat mass did not differ in mice based on the microbiota received

In this study, we first introduced the diet high in daidzein and formulated with fermentable fibers two weeks before colonization of the synthetic bacterial communities designed (Equol(-) and Equol(+) communities) in these gnotobiotic mice. Four weeks after colonization of the gut microbiota, we assessed the body weight, mesenteric fat mass, and gonadal fat mass at the time of euthanasia. Unsurprisingly, we did not discover any differences in these outcomes based on treatment groups (**Table 3**).

**Table 3.** Body weight and fat mass of male and female gnotobiotic mice colonized with the <u>non-equol producing (Equol(-)) and equol-producing (Equol(+)) communities.</u>.

|  | Males |  | Females |  |
| --- | --- | --- | --- | --- |
|  | Equol (-) | Equol (+) | Equol (-) | Equol (+) |
| Final body weight (g) | 27.19 ± 0.635 | 28.02 ± 0.845 | 22.04 ± 0.445 | 22.40 ± 0.271 |
| Mesenteric fat mass (g) | 0.28 ± 0.019 | 0.26 ± 0.030 | 0.18 ± 0.007 | 0.21 ± 0.016 |
| Gonadal fat mass (g) | 0.43 ± 0.028 | 0.45 ± 0.044 | 0.26 ± 0.019 | 0.29 ± 0.028 |

### The synthetic bacterial communities produce equol-producing capacity as designed in gnotobiotic mice fed a high daidzein diet

The synthetic communities designed include five phyla commonly found in human intestinal microbiomes, and the predominant phyla Bacteroidota and Bacillota are represented by a higher diversity of strains [42–45]. Before forming the communities, we tested each bacterial strain individually for its equol-producing capability *in vitro* and confirmed that equol production was only detected for the equol-producing strain of *A. equolifaciens*, as expected (**Table 4**). The equol-producing capability of these two synthetic communities was then tested *in vivo* by inoculating male and female germ-free mice to create gnotobiotic mice colonized with Equol(-) or Equol(+) communities. It has been suggested that females are more likely to become equol producers, so we included both sexes to observe potential sex differences [46]. Two weeks after colonization, the concentration of soy isoflavone daidzein and the microbial metabolite equol was assessed in the serum of gnotobiotic mice. Daidzein was present in all serum samples and did not differ significantly between groups within each sex (**Figure 1**). As expected, equol was absent in the serum of all mice colonized with the Equol(-) communities and was detected in the serum of all mice colonized with the Equol(+) communities. Sex differences were not observed as the levels of daidzein and equol did not differ between males and females statistically (data not shown).

**Table 4.** Equol-producing capacity of each bacterial strain cultured in vitro. Each bacterial isolate used in this study was grown in daidzein supplemented culture broth and the concentrations of daidzein and equol were analyzed using an LC-MS/MS analysis.

| Species | Strain | Daidzein (μM) | Equol (μM) |
| --- | --- | --- | --- |
| *Bacteroides caccae* | ATCC 43185 | 46.18 | 0 |
| *Bacteroides thetaiotaomicron* | DSM 2079 | 7.55 | 0 |
| *Bacteroides uniformis* | ATCC 8492 | 30.42 | 0 |
| *Roseburia intestinalis* | DSM 14610 | 5.07 | 0 |
| *Faecalibacterium duncaniae* | DSM 17677 | 44.43 | 0 |
| *Agathobacter rectalis* | DSM 17629 | 13.35 | 0 |
| *Coprococcus comes* | ATCC 27758 | 36.14 | 0 |
| *Akkermansia muciniphila* | DSM 22959 | 28.04 | 0 |
| *Providencia stuartii* | ATCC 25826 | 40.21 | 0 |
| *Collinsella aerofaciens* | ATCC 25986 | 34.81 | 0 |
| *Adlercreutzia equolifaciens* | DSM 19450 | 3.12 | 29.05 |

**Figure 1.**
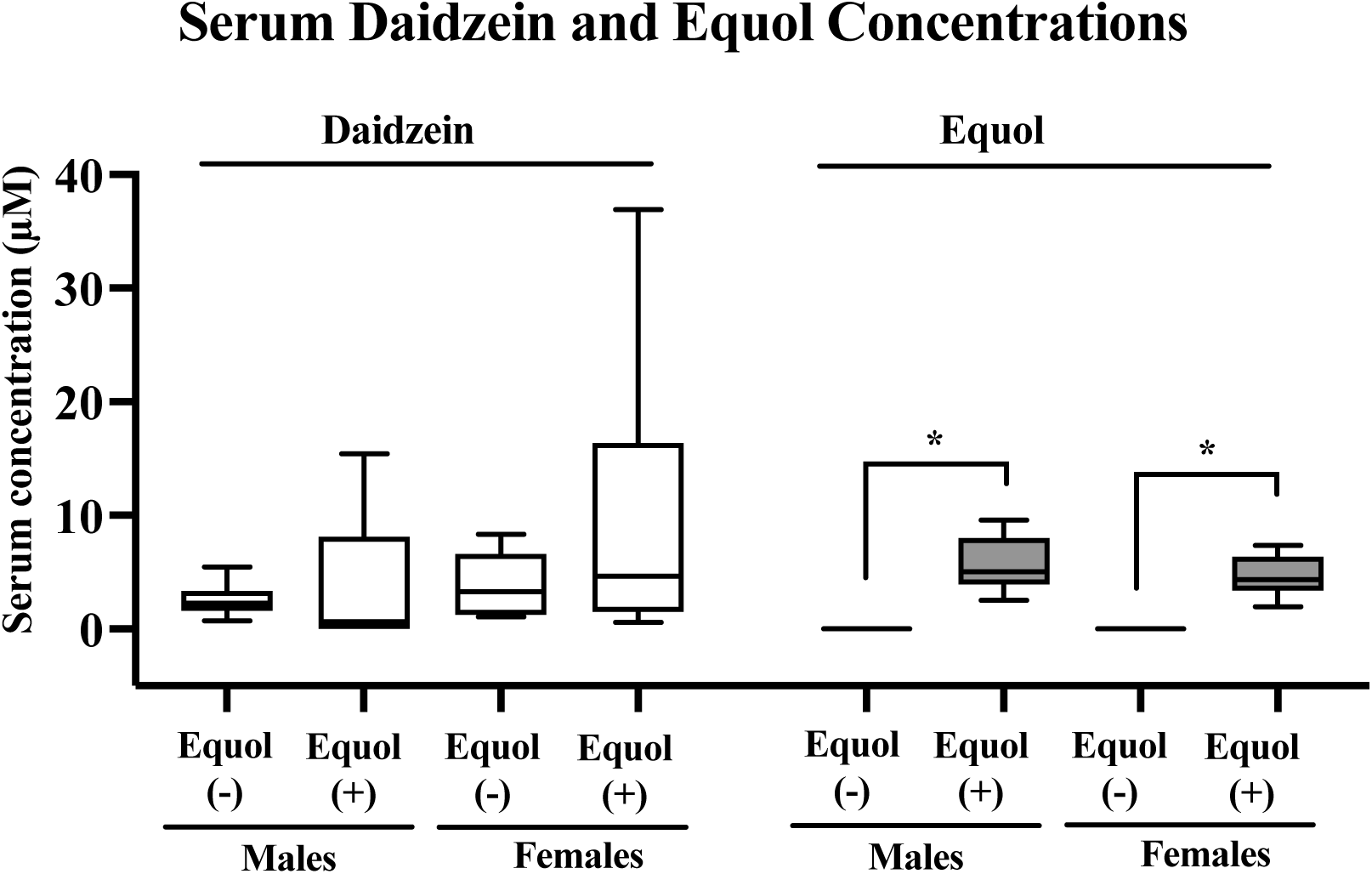
Concentrations of serum daidzein and equol in male and female gnotobiotic mice colonized with the non-equol producing (Equol(-)) and equol-producing (Equol(+)) communities. Box plots illustrate the median with a central line inside the box, which encompasses the interquartile range (IQR). Whiskers extend from the box to the lowest and highest data points. Statistical significance is denoted with asterisks.

### Equol-producing status did not affect the concentration of short-chain fatty acids in the cecum of gnotobiotic mice

Greater short-chain fatty acid productions have been suggested to support the synthesis of equol, likely through increasing hydrogen availability from fermentable substrates to provide electron donors in the bioconversion of daidzein to equol [32]. Therefore, we formulated the mouse diet to contain various fermentable fibers to not only facilitate the colonization of the synthetic bacterial communities, but also the synthesis of equol. However, to ensure that the presence of the equol-producing strain of *A. equolifaciens* will not alter SCFA, which may pose a confounding factor, we have assessed the level of SCFA in the cecal content of these gnotobiotic mice.

SCFA including acetic acid, propionic acid, butyric acid, isobutyric acid, valeric acid, and isovaleric acid were analyzed by a GC-MS. There were no significant differences in cecal SCFA between groups colonized with Equol(-) and Equol(+) communities within each sex (**Figure 2**). A significant difference in cecal propionic acid level was observed with male mice colonized with Equol(+) community having greater (p=0.02) propionic acid than the female mice colonized with the same microbial community.

**Figure 2.**
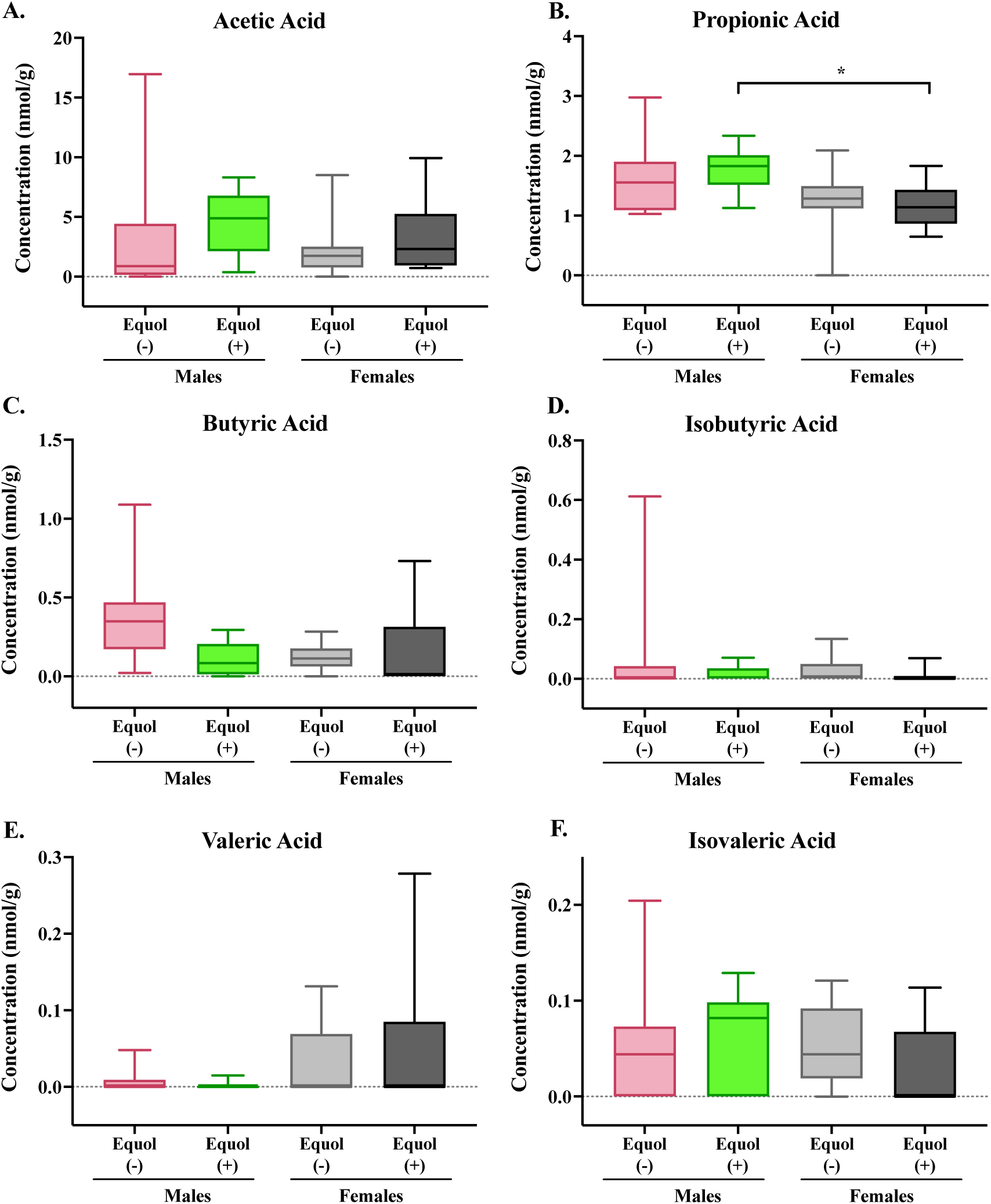
Concentrations of short-chain fatty acids in the cecal content of male and female gnotobiotic mice colonized with non-equol producing (Equol(-)) and equol-producing (Equol(+)) communities. Box plots illustrate the median with a central line inside the box, which encompasses the interquartile range (IQR). Whiskers extend from the box to the lowest and highest data points. Statistical significance is denoted with asterisks.

### Community-level assessment of the gut microbiota revealed differences between the bacterial inoculants and mouse gut, but similarity exists among treatment groups in mice

To assess the colonization similarity of each synthetic bacterial community designed in male and female germ-free mice, the community composition of the gut microbiota was assessed using 16S rRNA gene sequencing of the mouse cecal contents and bacterial inoculants used to colonize these mice. PCoA plots show that the mouse cecal microbiota clustered separately (p<0.05) from the inoculants, as expected (**Figure 3A**). For the bacterial inoculants, the Equol(-) and Equol(+) groups cluster tightly together without statistical differences. When assessing the mouse cecal microbiota alone, there were significant differences between Equol(+) females and Equol(-) males (**Figure 3B**; Bray-Curtis PERMANOVA, p= 0.008; Weighted UniFrac PERMANOVA, p= 0.016) and between Equol(+) males and Equol(-) males (Bray-Curtis PERMANOVA, p=0.002; Weighted UniFrac PERMANOVA, p=0.002). There were no significant differences in the dispersion of the data of the groups, indicating that the statistical significance observed was driven by differences in the mean distances between groups. However, a large overlap of the gut microbiota among the treatment groups suggests that the colonization between groups was generally similar whether accounting for the phylogenetic relationship between the bacterial species or not.

**Figure 3.**
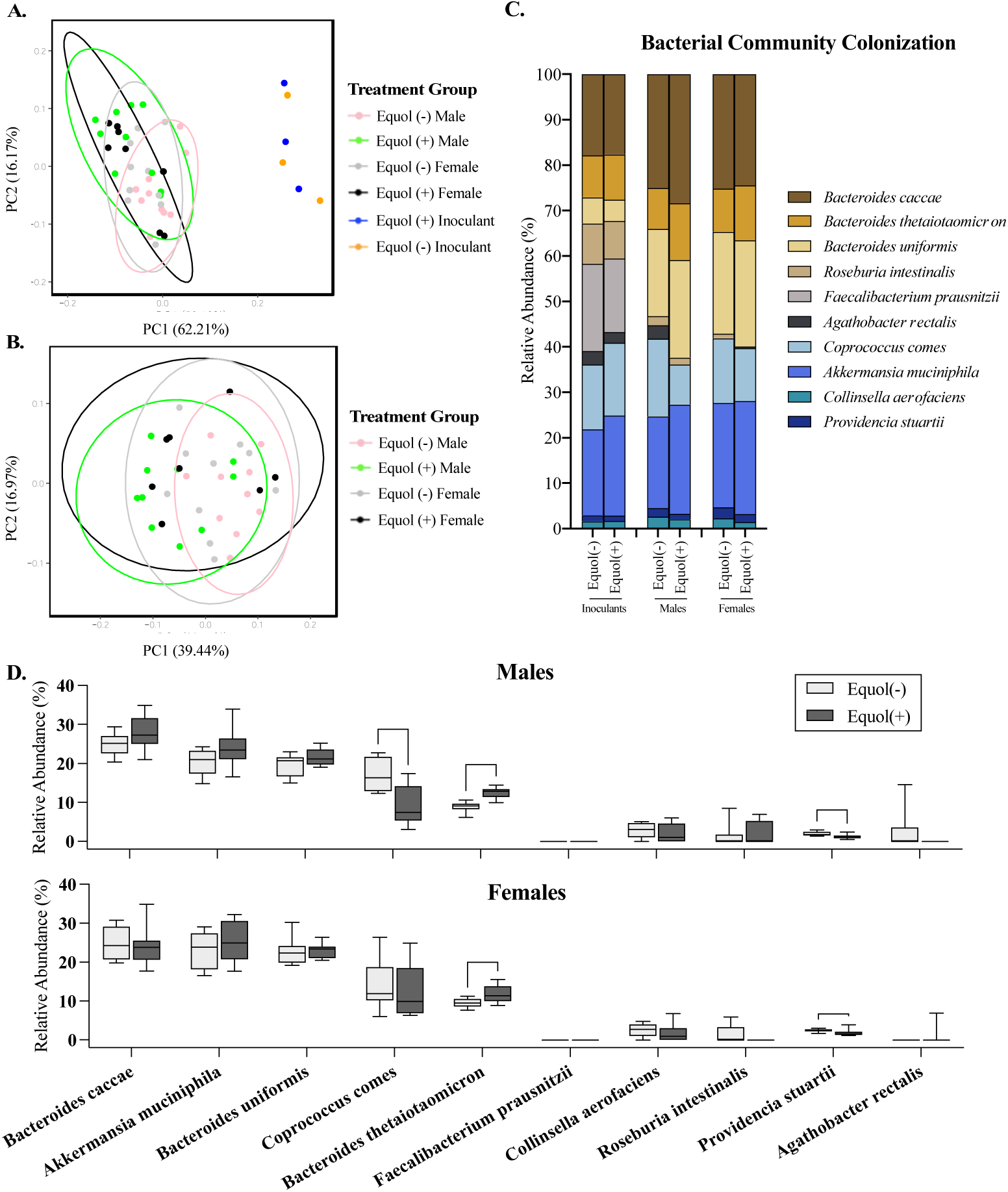
Microbiota profiling of the cecal microbiota in male and female gnotobiotic mice colonized with the non-equol producing (Equol(-)) and equol-producing (Equol(+)) communities using 16S rRNA sequencing. Community level differences among the inoculants of synthetic bacterial communities designed and those eventually colonized in the cecal content of mice was analyzed using the Bray-Curtis metric and displayed using a principal coordinate analysis (PCoA) plot (A). Analysis was also performed in the cecal microbiota of mice, without the inoculants (B). Each sphere in the PCoA plots represents a unique microbial community, with communities closer in proximity being more similar to each other. The percent of variation explained by each axis of the PCoA plot is listed in parentheses on each axis. The average relative abundance of each strain within a treatment group is summarized (C), with differential abundance of each taxon being compared between Equol(-) and Equol(+) in males and females (D). Statistical significance is denoted with asterisks.

### The relative abundance of some bacterial taxa in mice colonized with Equol(-) and Equol(+) differ significantly statistically, but are still similar proportionally

The mean relative abundance of the bacterial species in the inoculants and mouse cecal contents was assessed. While the inoculants were prepared by pooling equal OD_600_, except for the two strains that cannot be grown to OD_600_ of 1.0 (*A. equolifaciens* and *P. stuartii*), the relative abundance of each bacterial strain was not proportionally equal in the inoculants (**Figure 3C**). This is not surprising, considering the biases present in the nature of the methodology chosen for sequencing, but confirms that all but one taxon, *A. equolifaciens*, were present at a detectable range in the inoculants prepared. In the mouse cecal content, on the other hand, *A. equolifaciens and F. duncaniae* were not detectable in the cecal microbiota using the 16S rRNA sequencing method through the Illumina platform (**Figure 3C&D**). The relative abundance of *B. thetaiotaomicron* was greater (p<0.05) in both male and female mice colonized with the Equol(+) community compared to those colonized with the Equol(-) community (**Figure 3D**). The relative abundance of *P. stuartii* was lower (p<0.05) in both male and female mice colonized with the Equol(+) community than the Equol(-) community. And *C. comes* was present at a lower relative abundance in male mice colonized with Equol(+) community, but not in female mice. For the two strains that were not detected in the cecal microbiota of the mice using sequencing, we performed targeted qPCR to assess their presence. As expected, *A. equolifaciens* was present in the inoculant and mouse cecal microbiota of the Equol(+) community at 10^5^-10^6^ CFU and absent in samples associated with Equol(-) community (**Figure 4A**). *F. duncaniae*, on the other hand, was only detected in the inoculants but not the cecal content.

**Figure 4.**
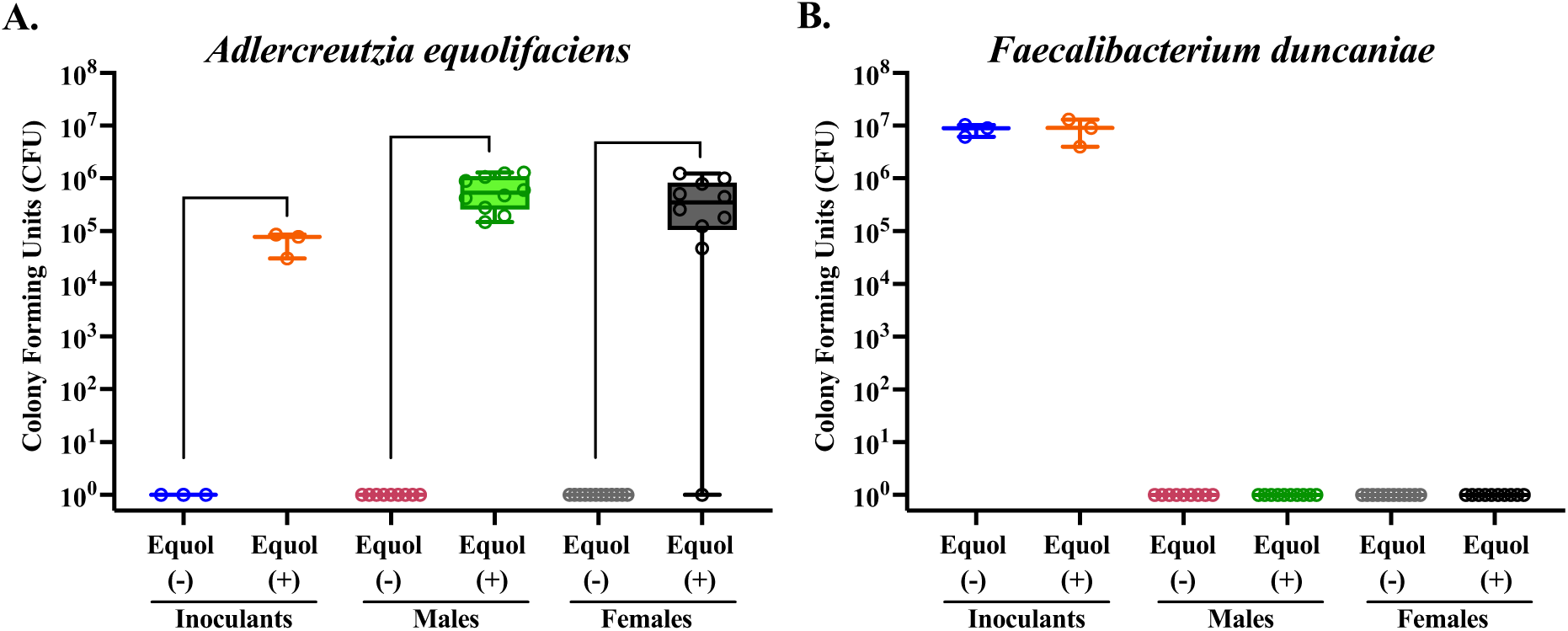
Quantification of *Adlercreutzia equolifaciens* and *Faecalibacterium duncaniae* strains in mouse cecal contents and inoculants using qPCR. Standard curves were generated for each strain to quantify the level of each strain present in the inoculants and mouse cecal samples for each treatment group. The average CFU/ml of each strain within each group was calculated. Statistical significance is indicated as follows: ∗p<0.05; ∗∗p < 0.01; ∗∗∗p < 0.001. Note: To ensure visibility on a logarithmic scale, values of 0 were plotted as 1, allowing all data points to be visible.

## Discussion

As most rodent models harboring natural microbiota are equol producers, outcomes generated from studies related to dietary soy and health are likely biased towards humans who are equol producers. The current study aimed to create a gnotobiotic mouse model without equol-producing capacity by combining non-equol-producing bacteria strains to form a synthetic community. An equol-producing bacterial strain was added to the non-equol-producing community to form a synthetic bacterial community with equol-producing capability. Analysis of blood equol concentration successfully demonstrated our ability to create gnotobiotic mice with divergent equol-producing capabilities that consistently display the expected equol-producing phenotypes. Even though strain-specific differences were detected between Equol(-) and Equol(+) communities in this study, variability of the communities remained low in the microbiomes of the equol producers and non-equol producers in this model compared to if using complex human microbiome samples to colonize the mice [19,25,42]. This mouse model will allow us to mechanistically test and isolate the effect of endogenous bacterial equol production on disease predisposition and prognosis related to soy and/or equol.

The average serum levels of equol observed in the gnotobiotic mouse model in the current study are similar to levels observed in previous soy studies of rodents (1.7-10µM) [24,47–49]. Interestingly, the concentrations of serum equol that we and others have found in rodents are much higher than those reported in humans. Equol-producing humans have been defined to be those with plasma equol concentrations of 83 nmol/L, or 0.083µM [50]. A human study showed improved cardiovascular health outcomes at average serum equol concentrations of 0.236µM [16]. These data suggest that, even when colonized with human-derived bacterial strains, rodents are superior equol-producers that produce 7-120 times higher levels of equol than humans. However, the lack of reporting on serum equol levels in numerous soy-related human studies complicates the assessment of whether the high serum concentrations of equol observed in this rodent model are attainable in humans.

A few bacterial genera, particularly those within the *Eggerthellaceae* family, have been identified to have the equol-producing capacity, including *Adlercreutzia, Slackia*, *Eggerthella*, *Paraeggerthella, Asaccharobacter,* and *Enterorhabdus,* as summarized in a narrative review paper that was published by our research team [51]. Some genera outside of the *Eggerthellaceae* family have been identified as well, including certain species and strains within genera *Lactobacillus, Lactococcus,* and *Bifidobacterium* [12,52–56]. In this study, we chose to use *A. equolifaciens* as the equol-producing strain. We first utilized the 16S rRNA sequencing technique to assess the microbial communities that were used to inoculate mice and also those eventual colonizers in the cecal content of mice. Although an average sequencing depth of 19,182 reads per sample was achieved, indicating sufficient coverage, equol-producing *A. equolifaciens* strain were not detected in any of the mice in this study through 16S rRNA sequencing even though equol was detected in the serum. This is somewhat surprising as approximately 4% of the total sequencing reads have been reported to be *A. equolifaciens* in a recent report assessing equol-producing strains in human microbiota using a similar technique [57]. We speculated that a similar or even higher proportion of the total bacterial reads would be detected in a simplified synthetic bacterial community like those used in the current study. We then successfully confirmed the presence of *A. equolifaciens* in our study at an approximation of 10^6^ CFU using qPCR. Unfortunately, the one published study utilizing gnotobiotic mice to create divergent equol phenotype did not sequence or quantify equol-producing bacterial strain and did not assess the community structure of the gut microbiota so we are unable to compare the colonization in gnotobiotic rodent models [23]. Primer bias may contribute to reads associated with *A. equolifaciens* being lower than detectable in 16S rRNA-targeted gene sequencing. Future attempts can be made using different sequencing methods, such as short-read or long-read metagenomic sequencing to detect *A. equolifaciens* in this synthetic community. Our study emphasizes that the sequencing approach targeting the 16S rRNA gene may not be sensitive enough to detect the presence of bacterial strains with important microbial phenotypes, such as equol production.

One of the strains included in our synthetic bacterial community, *Faecalibacterium duncaniae*, was recently proposed to be renamed as such [58]. *F. duncaniae* used to be classified as *Faecalibacterium prausnitzii* and it is a butyrate-producing bacterium and a very strict anaerobe [59]. One limitation of this study is that *F. duncaniae* was not able to colonize, despite being present in the inoculant. The presence of oxygen in the stomach and small intestine of germ-free mice due to the complete lack of oxygen may have hindered the colonization of this strain. Others have had some success in improving the colonization of synthetic communities by consecutive inoculation [60,61]. We have attempted similar consecutive inoculation in a separate study by providing a second oral gavage a week after the first inoculation took place. However, this effort did not significantly affect the strain colonization of *F. duncaniae*. It is worth noting that another butyrate-producing strain is present in this community that was successfully colonized. Nevertheless, a replacement of *F. duncaniae* could be considered if the presence of an F. *prausnitzii*-like strain is desired.

## Conclusions

The gnotobiotic mouse model system designed in this study can successfully establish microbiomes with divergent equol-producing phenotypes *in vivo* and be utilized to establish a causal relationship between equol and the consumption of soy isoflavones. Breeding of these gnotobiotic mice with disparate equol phenotypes would be possible to assess developmental impact of equol with potential transgenerational influence from parents with the same equol phenotype. This model system provides great potential for future research on the health benefits of equol, as it has far fewer confounding variables than human studies. The benefit of having a true negative control for equol production sets this model apart from conventional rodent models, and the minimization of differences between the microbiomes of the mice further reduces confounding variables. The use of this system can provide causative evidence linking the benefits of soy consumption to the microbial production of equol. The applications of this model system are broad, as this model can be used as a foundation to study a range of health outcomes in future studies such as a reduced risk of breast cancer, improved cardiovascular health, and improved bone health. Further, it also can be used to determine potential health concerns regarding the consumption of soy, such as the reproductive health of men.

## Author Contributions

Conceptualization, T.W.C. and L.R.; validation, L.L., A.S., and T.W.C.; formal analysis, L.L. and A.S.; investigation, L.L. and A.S.; data curation, L.L., A.S., and S.L.; writing—original draft preparation, L.L., A.S., and T.W.C.; writing—review and editing, T.W.C. and L.R..; visualization, L.L., A.S., and T.W.C.; supervision, T.W.C.; funding acquisition, T.W.C. All authors have read and agreed to the published version of the manuscript.

## Institutional Review Board Statement

The animal study protocol was approved by the Purdue Animal Care and Use Committee.

## Acknowledgments

We would like to thank the animal care staff at the Purdue Gnotobiotic Facility for their assistance with our rodent work, and acknowledge the Purdue Genomics Core at the Purdue Bindley Bioscience Center.

---

**Supplemental Table 1.** Bacterial culture media and incubation times. The following broth and agar were used for the growth of each strain in this study. All strains were grown anaerobically at 37°C. Agar incubation requires an additional 1-2 days to achieve optimal colony growth.

| Species | Culture Broth | Agar | Broth Incubation Time |
| --- | --- | --- | --- |
| *Bacteroides caccae* | Anaerobic Basal Broth (ABB, DSMZ Medium 1203a) | Brucella Blood Agar (BBA, Thermo Scientific (Cat. No. BD 211086) with 5% blood) | 12-24 hours |
| *Bacteroides thetaiotaomicron* | ABB | BBA | 12-24 hours |
| *Bacteroides uniformis* | ABB | BBA | 36-48 hours |
| *Roseburia intestinalis* | Yeast Casitone Fatty Acid (YCFA) broth (see Table S2) | YCFA agar | 12-24 hours |
| *Faecalibacterium duncaniae* | Reinforced Clostridia Media (RCM, Thermo Scientific (Cat. No. CM0149B)) broth | RCM agar | 12-24 hours |
| *Agathobacter rectalis* | PGY (DSMZ Medium 104) | BBA | 36-48 hours |
| *Coprococcus comes* | ABB | BBA | 12-24 hours |
| *Akkermansia muciniphila* | Anaerobic Brain Heart Infusion broth (A-BHI, ATCC Medium 2187) + 0.4% Mucin | Tryptic Soy Agar (TSA) + 5% Blood (ATCC Medium 260) | 36-48 hours |
| *Providencia stuartii* | Nutrient Broth (NB, ATCC Medium 3) | TSA + 5% Blood | 12-24 hours |
| *Collinsella aerofaciens* | ABB | BBA or RCM | 12-24 hours |
| *Adlercreutzia equolifaciens* | Wilkins-Chagrin (WC, Thermo Scientific (Cat. No. CM0643B)) broth | WC agar | 48-72 hours |

